## Supplementary Figures for "The olfactory receptor SNIF-1 mediates foraging for leucine-rich diets in *C. elegans*"

### Supplemental Information

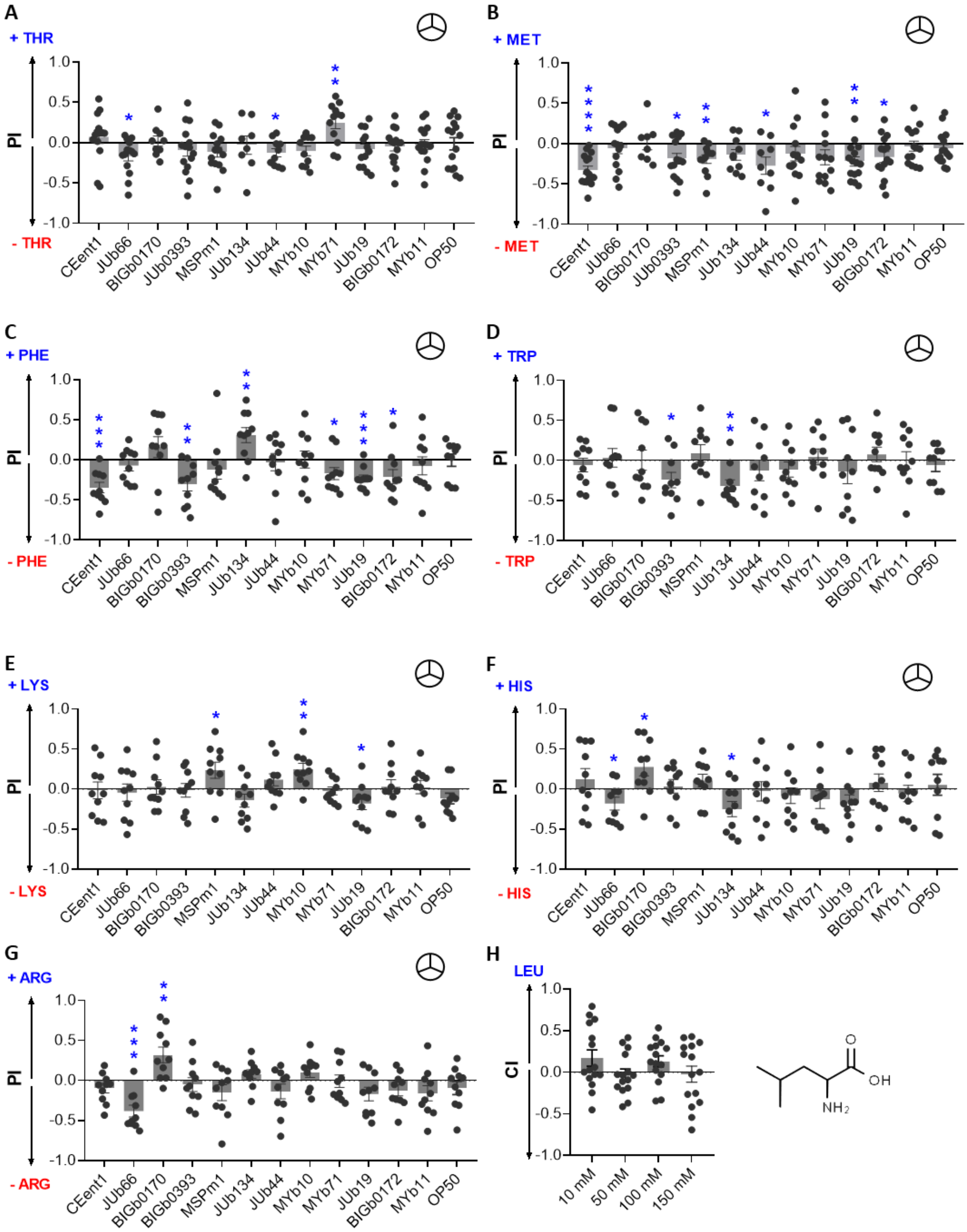


**Figure S1. *C. elegans* does not prefer some EAA supplemented diets**

Preference index (PI) of WT worms in an 'odor-only' diet preference assay for individual diet supplemented with (A) 5 mM threonine (+ THR), (B) 5 mM methionine (+ MET), (C) 5 mM phenylalanine (+ PHE), (D) 5 mM tryptophan (+ TRP), (E) 5 mM lysine (+ LYS), (F) 5 mM histidine (+ HIS) and (G) 5 mM arginine (+

ARG). ☮ symbol indicates ‘odor-only’ preference assays. Significant differences are indicated as * $P \leq 0.05$, ** $P \leq 0.01$, and **** $P \leq 0.0001$ determined by one-sample *t*-test.
(H) Chemotaxis index (CI) for WT worms to leucine (LEU) at various concentrations.
Error bars indicate SEM ($n \geq 10$).

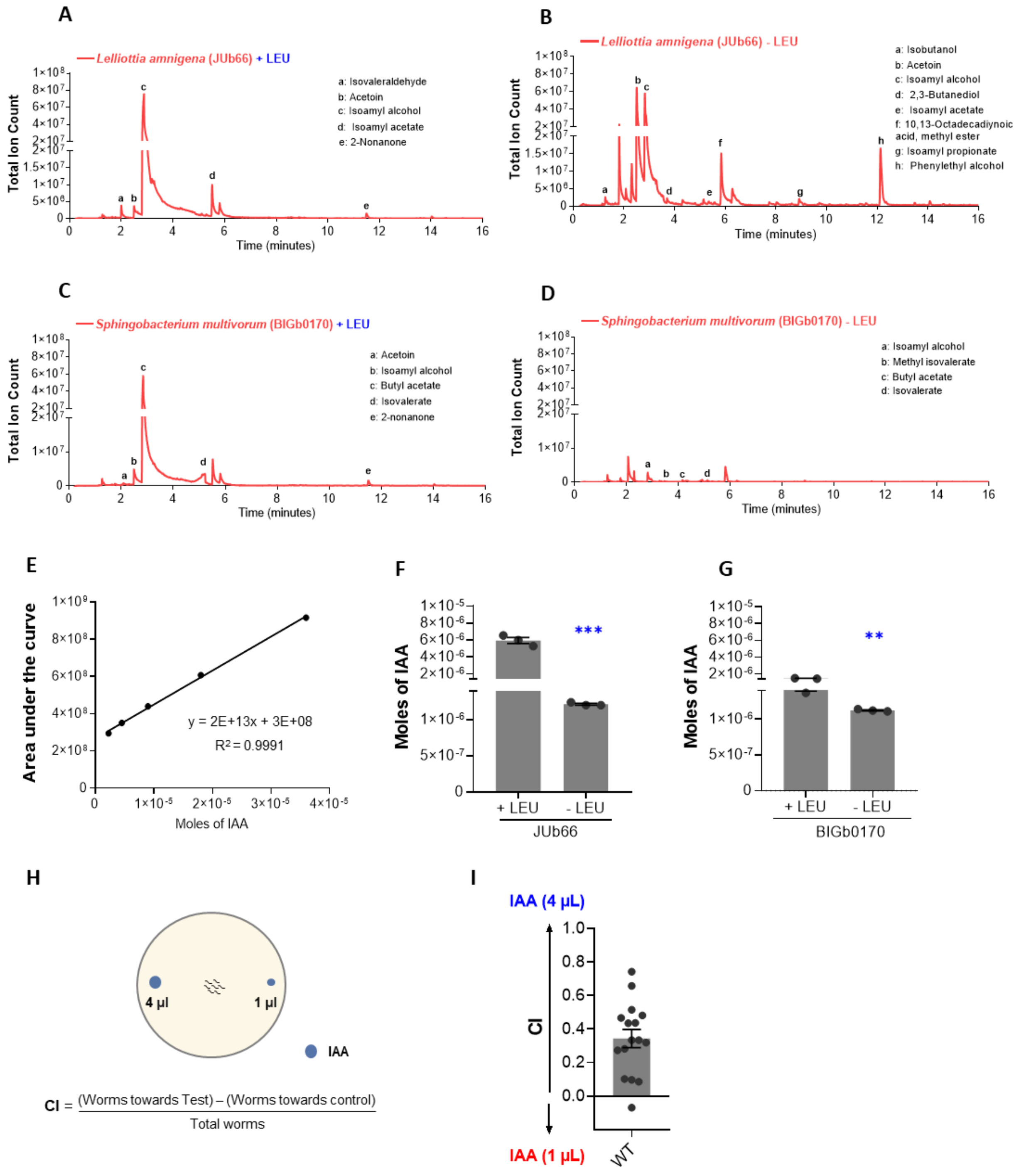


**Figure S2. Increased level of isoamyl alcohol upon leucine-enriched influence worms' diet preference**

GC-MS/MS profile of odors produced by JUb66 in (A) leucine supplemented (+ LEU) and (B) leucine unsupplemented (- LEU) conditions.

GC-MS/MS profile of odors produced by BIGb0170 in (C) leucine supplemented (+ LEU) and (D) leucine unsupplemented (- LEU) conditions. Unmarked peaks represent masses contributed by fiber or media alone (n≥3).

(E) Standard curve showing the correlation between the area under the curve for odor peak and the moles of IAA.
Absolute abundance of IAA (in moles) produced by (F) JUb66 and (G) BIGb0170 under leucine supplemented and unsupplemented conditions. ** $P \leq 0.01$, *** $P \leq 0.001$ as determined by the two-tailed unpaired *t*-test. Error bars indicate SEM (n≥3).
(H) Schematic representation of chemotaxis assay between different concentrations of IAA.
(I) Chemotaxis index (CI) of WT worms to IAA (4μL over 1μL). Error bars indicate SEM (n≥15).

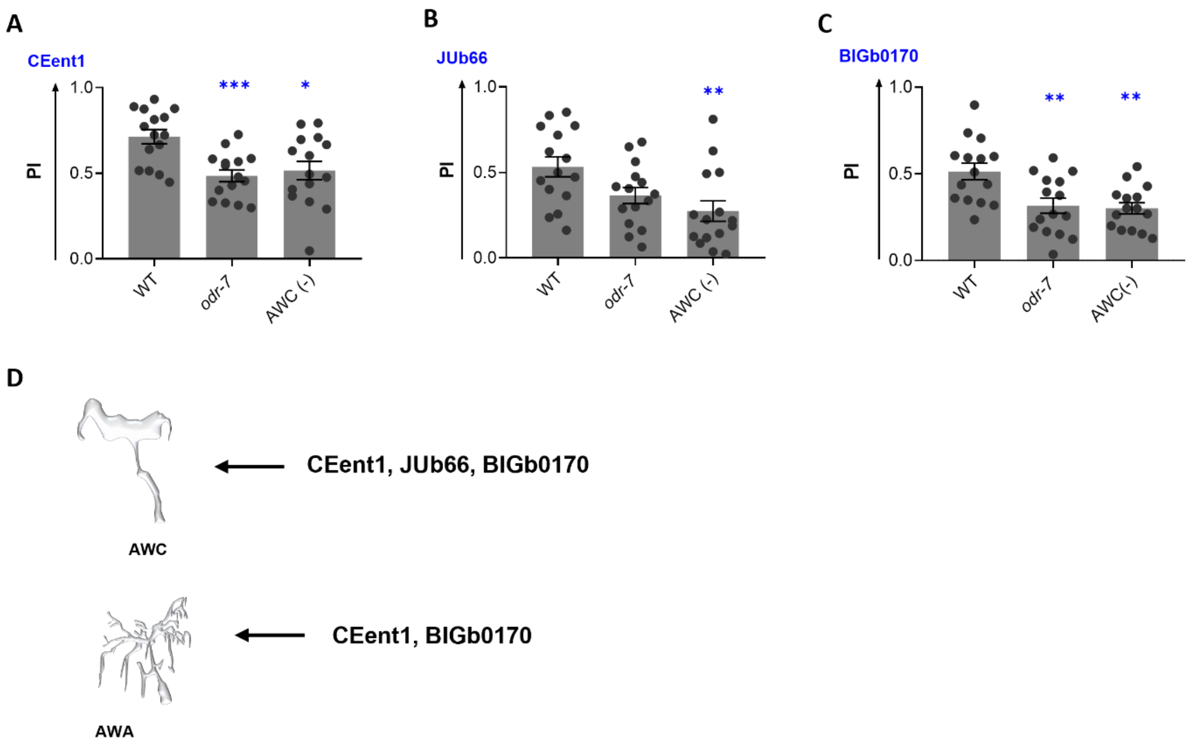


**Figure S3. AWA and AWC neurons facilitate the diet preference of *C. elegans***
Preference index (PI) of WT, *odr-7*, and AWC(-) worms in a diet preference assay for (A) CEent1, (B) JUb66, and (C) BIGb0170 over *E. coli* OP50.
(D) Summary representing the role of AWA and AWC odor sensory neurons in diet preference.
Significant differences are indicated as * P ≤ 0.05, ** P ≤ 0.01, and *** P ≤ 0.001 determined by one-way ANOVA followed by post hoc Dunnett’s multiple comparison test. Error bars indicate SEM (n=15).

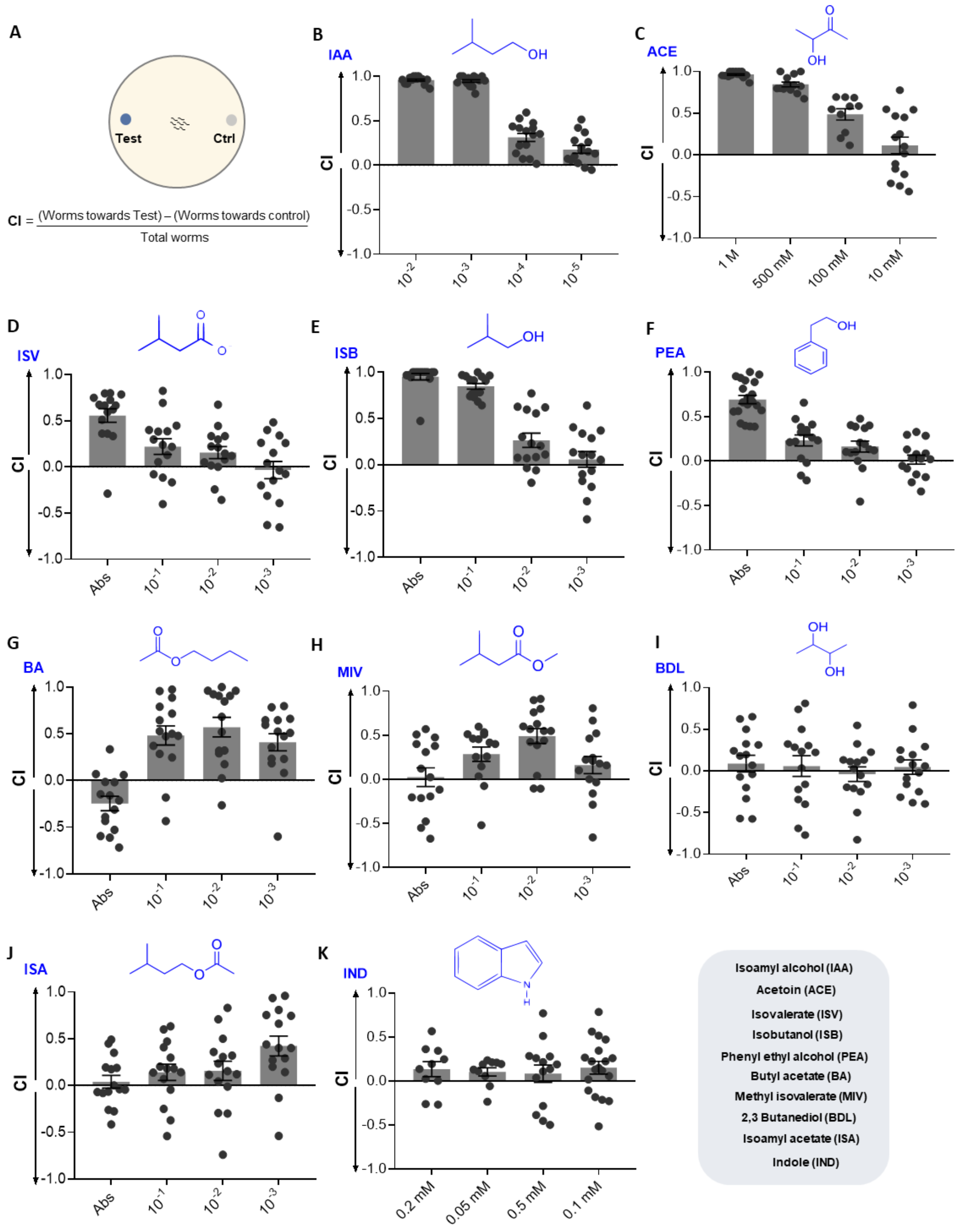


**Figure S4. Dose-dependent chemotaxis response of *C. elegans* to odors produced by preferred bacteria**

(A) Schematic representation of *C. elegans*' chemotaxis assay.

Chemotaxis index (CI) for WT worms to (B) isoamyl alcohol (IAA), (C) acetoin (ACE), (D) isovalerate (ISV), (E) isobutanol (ISB), (F) phenylethyl alcohol (PEA), (G) butyl acetate (BA), (H) methyl isovalerate (MIV), (I)

2,3-butanediol (BDL), (J) isoamyl acetate (ISA), and (K) indole (IND) to various concentrations of chemicals. Error bars indicate SEM (n=15).

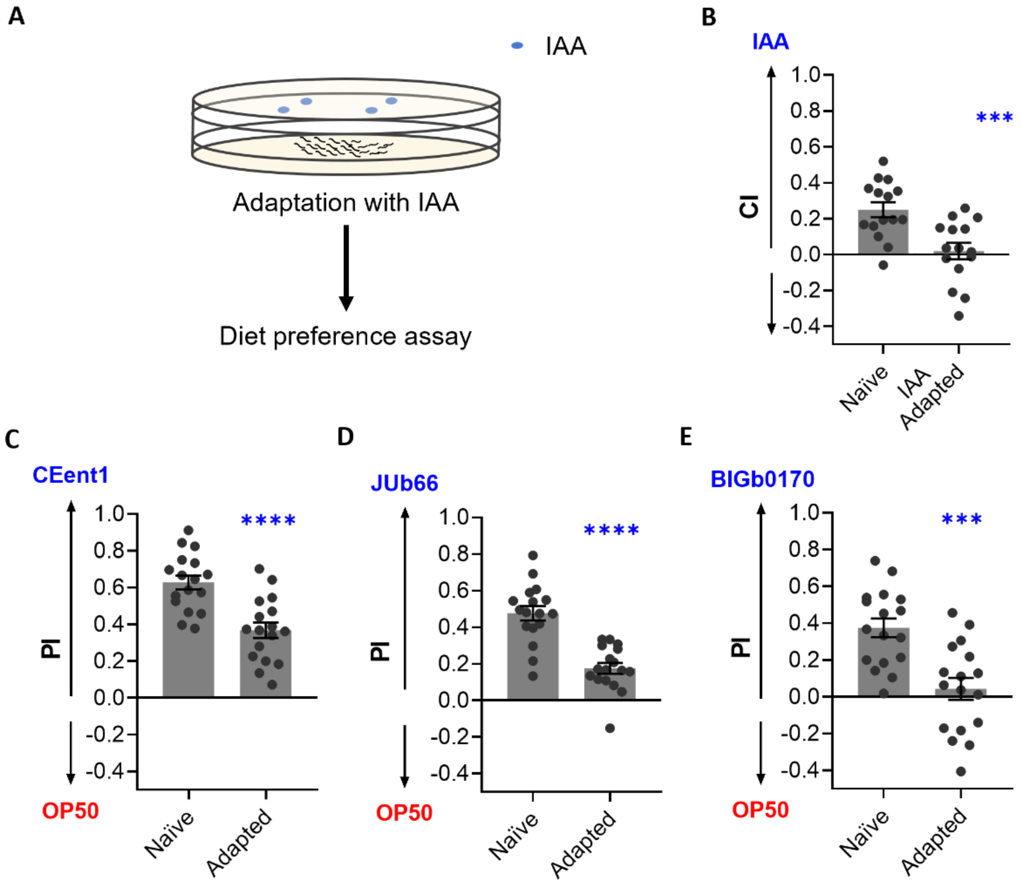


**Figure S5. *C. elegans* utilizes isoamyl alcohol to make dietary preferences**

(A) Schematic representation of the odor adaptation regimen followed by diet preference assays.

(B) Chemotaxis index (CI) to IAA for naïve worms and worms adapted with IAA.

Preference index (PI) of naïve worms and worms adapted with IAA in a diet preference assay for (C) CEent1, (D) JUb66, and (E) BIGb0170 over *E. coli* OP50.

Significant differences are indicated as ** P ≤ 0.01, *** P ≤ 0.001, and **** P ≤ 0.0001 determined by one-way ANOVA followed by post hoc Dunnett's multiple comparison test. Error bars indicate SEM (n=15).

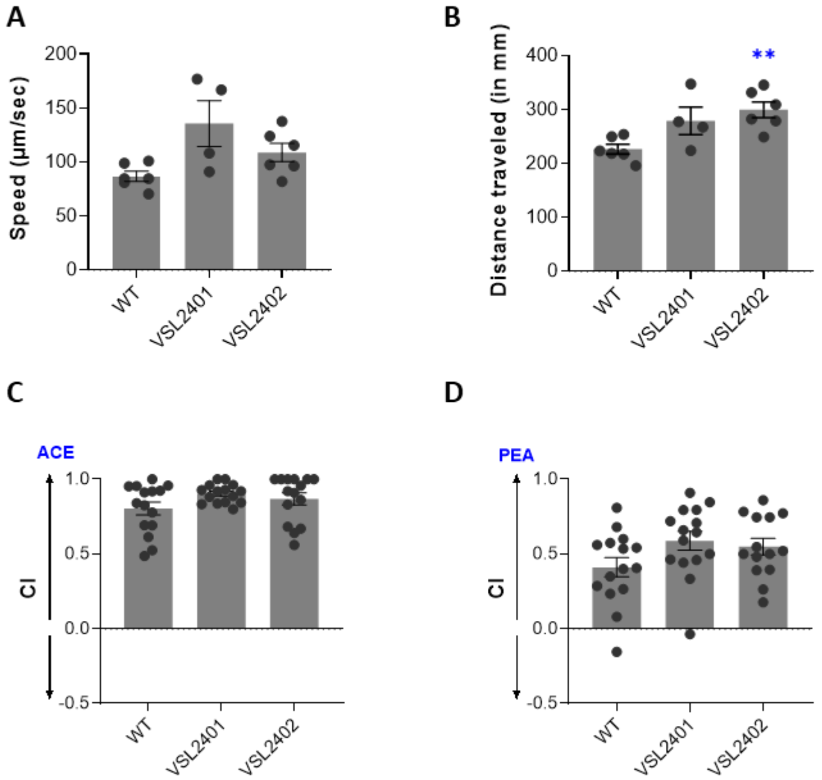


**Figure S6. Differential effect of SNIF-1 on locomotion of *C. elegans***

(A) Average speed of worms (μm/ second), and (B) total distance traveled (in mm) by WT, VSL2401, and VSL2402 worms in a chemotaxis arena in response to IAA in 30 minutes (n≥3).

(B) Chemotaxis index (CI) of WT, VSL2401, and VSL2402 worms for (C) 500 mM acetoin (ACE), and (D) absolute phenylethyl alcohol (PEA). (n≥15).

Significant differences are indicated as ** P ≤ 0.01 determined by one-way ANOVA followed by post hoc Dunnett's multiple comparison test. Error bars indicate SEM.
